# HipHap: Haplotype Assignment and Confidence Scoring for Diploid Reference Genomes

**DOI:** 10.64898/2026.09.24.754153

**Authors:** Jakob M. Heinz, Maximillian G. Marin, Matthew Meyerson, Heng Li

**Affiliations:** Department of Biomedical Informatics, Harvard Medical School, Boston, MA, USA; Department of Data Science, Dana-Farber Cancer Institute, Boston, MA, USA; Cancer Program, Broad Institute of MIT and Harvard, Cambridge, MA, USA; Department of Medical Oncology, Dana-Farber Cancer Institute, Boston, MA, USA; Department of Genetics, Harvard Medical School, Boston, MA, USA

**Keywords:** Diploid Assembly, Alignment, Phasing, Long-read

## Abstract

Diploid genome assemblies are now routinely available, but most read aligners were designed for haploid references, which have long been the gold standard. When reads are aligned to a diploid assembly, the aligner sees two nearly identical alignments to either haplotype, thus reducing the mapping quality (MapQ) score to reflect this ambiguity. This can cause downstream tools to discard reads from easily mappable regions. Here, we present HipHap (HIgh-Performance HAPlotype assigner) to resolve this issue by aligning reads to each haplotype assembly separately and assigning each read to its best-supported haplotype. HipHap increased the fraction of reads mapping to the diploid genome with high MapQ, outperforming alignment to either parental haplotype. Additionally, we introduce a haplotype assignment quality score (HapQ) in HipHap to quantify confidence in the haplotype of origin of a read. HipHap is implemented in Rust, supports SAM, BAM, CRAM, and PAF formats, and is freely available at: https://github.com/jheinz27/hiphap.

## 1. Introduction

Diploid organisms, such as humans, have two homologous copies of each autosomal chromosome. However, genomic analyses have historically relied on a haploid reference genome (International Human Genome Sequencing Consortium, 2001). In haploid (or “primary”) assemblies, haplotypic variation is collapsed into a consensus sequence for each autosome to maximize contiguity (Garg et al., 2018). As a result, the current human reference genome, GRCh38, is a mosaic not only of individuals, but also of haplotypes, frequently switching between phases (Schneider et al., 2017; Liao et al., 2023). Until recently, using a haploid reference genome was a practical necessity because short read lengths and high sequencing error rates made accurate haplotype assembly extremely difficult. However, developments in high-accuracy long-read sequencing enabled the Telomere-to-Telomere (T2T) consortium to complete the first gap-free human genome assembly in 2022 (Nurk et al., 2022). The consortium circumvented the challenge of haplotype phasing in assembly by using a functionally haploid cell line, CHM13. Now, with high-accuracy long reads and further advances in assembly algorithms, assemblers such as Verkko and Hifiasm (Rautiainen et al., 2023; Cheng et al., 2021, 2026) can routinely generate complete or near-complete haplotype-resolved diploid genome assemblies. A notable example is the Q100 HG002 assembly, which serves as a high-accuracy diploid reference assembly (Hansen et al., 2025).

A diploid reference genome provides a more accurate representation of the human genome. Yet, it also introduces new challenges, as many current bioinformatics tools were designed with the assumption of a haploid reference genome. For example, current aligners, such as Minimap2 (Li, 2018), assign mapping quality (MapQ) scores based on the confidence that a read is correctly aligned to the genomic locus it originated from. However, when reads are mapped to a diploid reference assembly, homologous loci on two haplotypes yield nearly identical alignments. The aligner interprets this as an ambiguous alignment, which artificially lowers the MapQ score. Variant callers and other downstream tools routinely filter on MapQ scores (Smolka et al., 2024; Zheng et al., 2025; Keskus et al., 2026), meaning this artificial reduction can lead to the exclusion of reads from regions that would be easily mappable to a haploid reference.

We present a “HIgh-Performance HAPlotype assigner”, or HipHap, to resolve the mapping ambiguity introduced by diploid reference genomes. Our method compares independent alignments of the same set of reads to each respective haplotype assembly and assigns each read to its best-supported haplotype. HipHap builds on a rudimentary preprocessing coding script in our recent Breakinator package (Heinz et al., 2026). HipHap is intended for long-read sequences, both genomic and transcriptomic, with sufficient length to permit haplotype assignment. Additionally, we introduce a haplotype assignment quality score (HapQ) to quantify confidence in the inferred haplotype of origin of a read. We propose using HapQ as an additional confidence measure alongside MapQ. While MapQ reflects confidence in the genomic position of an alignment (Li et al., 2009), HapQ reflects confidence in the haplotype of origin. We find that together the two scores provide a more comprehensive picture of alignment reliability with respect to a diploid reference genome.

## 2. Implementation

HipHap is implemented in Rust and takes as input two alignment files produced by aligning the same read set independently to each haplotype. To avoid loading entire files into memory, both files must be sorted by read name, which is the Minimap2 (Li, 2018) default output, so that records for the same read can be consumed in lockstep. HipHap uses the rust-htslib library (Bonfield et al., 2021; Köster, 2016) to read SAM/BAM/CRAM and PAF input files. It outputs summary statistics on the number of reads assigned to each haplotype, along with a merged alignment file that retains each read’s alignment to its better-supported haplotype, annotated with the calculated HapQ score (Section 2.3). The output alignment file can be sorted and coordinate-indexed with samtools (Li et al., 2009) or passed to downstream tools. Alternatively, HipHap can write partitioned output files, one for each haplotype, using the `-p` option.

### 2.1. Haplotype assignment

As summarized in Algorithm 1, for each read, HipHap groups primary, supplementary, and secondary alignment records into a cluster per haplotype alignment (Algorithm 1, lines 1 to 3). The whole cluster is written to the chosen output alignment file. Reads that are unmapped in both haplotypes can be discarded or written to the output file (user-specified) (Algorithm 1, lines 4 to 5). Reads that map to only one haplotype are assigned directly to that haplotype (Algorithm 1, lines 6 to 7). For reads that map to both haplotypes, a weighted alignment score is computed for each cluster (see Section 2.2), and the read is assigned to the haplotype with the higher score (Algorithm 1, lines 8 to 15). When both scores are equal, the haplotype assignment file is randomly chosen using the last bit of a deterministic hash (`twox hash::XxHash64`) of the read name to ensure reproducibility across runs (Algorithm 1, lines 16 to 17). Reads whose winning alignments span different chromosomes are output in a separate FASTA file, flagging them as potentially representing between-haplotype events (not represented in Algorithm 1 summary).

### 2.2. Weighted alignment score

When a read has a single end-to-end primary alignment to both haplotypes, comparing the two alignments is a straightforward comparison of their alignment scores (AS). However, when a read alignment is split into primary and supplementary records, a weighted approach is needed because the aligned intervals on the read may overlap, inflating the AS sum. For example, consider a 7,000 bp read, where the primary alignment covers bases 0– 5000, and the supplementary alignment covers bases 4000–7000. The 1,000 bases in the overlap (4000–5000) contribute to the alignment scores of both records, so summing alignment scores directly would double-count these shared bases and inflate the score of a split alignment.

#### Algorithm 1

HipHap: Haplotype-specific read assignment

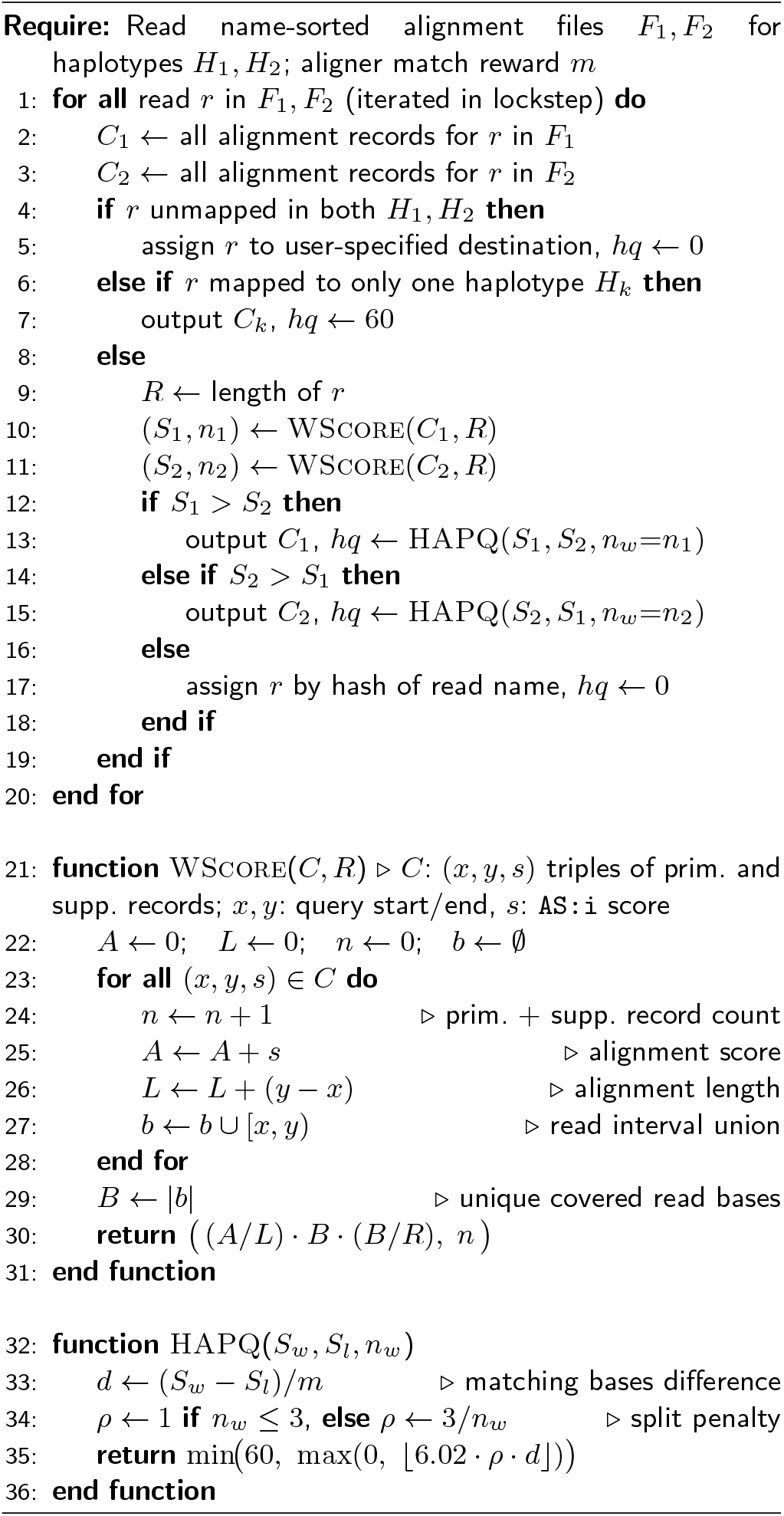

To correct for this inflation, we define a weighted alignment score *S* for each cluster of primary and supplementary records (Algorithm 1, lines 21–31). Secondary alignments are excluded from scoring but are retained in the output. For a cluster with *n* primary and supplementary records, let *a*_*i*_ denote the alignment score (`AS:i` tag) and *ℓ*_*i*_ the aligned query length of record *i*. The ratio *A/L* = *a*_*i*_*/ ∑ℓ*_*i*_ gives the average alignment score per aligned base of the read. Let *I*_*i*_ denote the interval on the read covered by record *i*, and let *B* = | ⋃ _*i*_ *I*_*i*_| be the number of unique read bases covered by any alignment after merging overlapping intervals. Finally, let *R* be the full read length. The weighted score is:

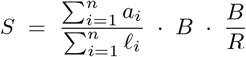

The first term (*A/L*) captures the average alignment quality per base, which is then scaled by *B* to give a total score over all uniquely aligned bases. The final term (*B/R*) is a coverage term that penalizes reads that are only partially aligned. This is needed when one haplotype contains a large deletion, as the aligner will only map the flanking regions, leaving the per-base quality comparable but much of the read unexplained. Consider a 10 kb read where haplotype 1 has a deletion, so only 5 kb aligns at an average score of 2.0 per base, while haplotype 2 aligns the full read at 1.0 per base. The alignment scores are identical (2.0 *×* 5000 = 1.0 *×* 10,000 = 10,000), but the *B/R* term breaks the tie: 2.0 *×* 5000 *×* 0.5 = 5,000 versus 1.0 *×* 10,000 *×* 1.0 = 10,000, favoring the more complete alignment despite its lower average score. The function also returns *n*, the count of primary and supplementary records in the cluster, which is used downstream by the HapQ split penalty (Section 2.3) as *n*_*w*_ (the count from the winning haplotype).

### 2.3. Calculating HapQ score

We introduce a haplotype assignment quality score (HapQ) to quantify the confidence that a read has been assigned to the correct haplotype, similarly to how MapQ quantifies the confidence that a read is correctly aligned to the position in the genome it originated from. The full HapQ derivation is shown in Appendix A. The probability of read sequence *r* of length *ℓ* being generated from a haplotype *h* without gaps is

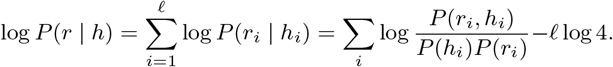

An aligner’s scoring matrix *s*(*a, b*) is proportional to the per-position log-odds ratio, for some scaling factor *c* > 0 (Durbin et al., 1998), therefore the per-position sum equals *S*(*r, h*)*/c*, where *S*(*r, h*) = *Σ* _*i*_ *s*(*r*_*i*_, *h*_*i*_) is the gapless alignment score (AS) between read *r* and haplotype *h*. Hence

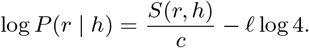

Given that we observe the read and not the haplotype, we seek to calculate the posterior probability that the higher-scoring (winning) haplotype *h*^(*w*)^ is the true source of read *r*, rather than the alternative (losing) *h*^(*l*)^, which is

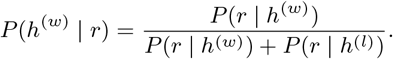

We report this confidence on the Phred scale as the probability that the winning haplotype assignment is wrong,

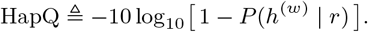

It remains to fix *c*. Note that under low error rate *ϵ*, match score *s*(*a, a*) *≈ c* log 4. Therefore

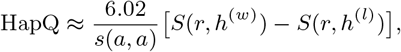

where *s*(*a, a*) is the match score used by the alignment software, and the constant 6.02 is the coefficient 10 log 4*/* log 10.

Similarly to BWA-MEM (Li, 2013), which applies several empirical adjustments to its raw mapping-quality calculation for sequence identity, alignment length, repeat content, and the number of near-optimal alternatives, we apply a single adjustment to the HapQ score reported by HipHap. We introduce a split-alignment penalty *ρ*, as a split alignment often reflects a structural variant that can introduce alignment artifacts, thereby reducing confidence in the correct assignment. Let *n*_*w*_ be the number of supplementary alignments supporting the winning haplotype. Because *n*_*w*_ can reach three when a true inversion is fully contained within a read, we penalize only *n*_*w*_ *>* 3:

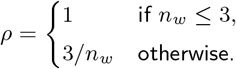

The remaining corrections that would be made by BWA-MEM are unnecessary here because we compare two alignments of the same read against two nearly identical haplotypes, so sequence identity, read length, and repeat content are similar between the competing alignments and cancel in the score difference *S*(*r, h*^(*w*)^) *− S*(*r, h*^(*l*)^). The final HapQ calculation used by HipHap is then

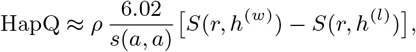

clamped to the range [0, 60]. This calculation corresponds to the HAPQ function in Algorithm 1, lines 32–36. The HapQ framework can be extended to gap penalties if they are on a similar scale as mismatches.

## 3. Results

We compared the MapQ distribution of reads using HipHap relative to two commonly used strategies: aligning reads to a single haplotype or to the full diploid assembly. We used publicly available PacBio HiFi reads (Pacific Biosciences, 2026) from the well-characterized HG002 cell line, as it has a high-quality diploid reference assembly available (Hansen et al., 2025). Only primary alignments to autosomes were considered for all following analyses. As shown in Figure 1a, aligning these reads to the entire diploid reference genome resulted in only 12.2% of reads mapping with a MapQ score of 60. HipHap restored this to 95.6%, an additional improvement over the individual maternal (94.8%) and paternal (93.9%) haploid alignments.

**Figure 1.**
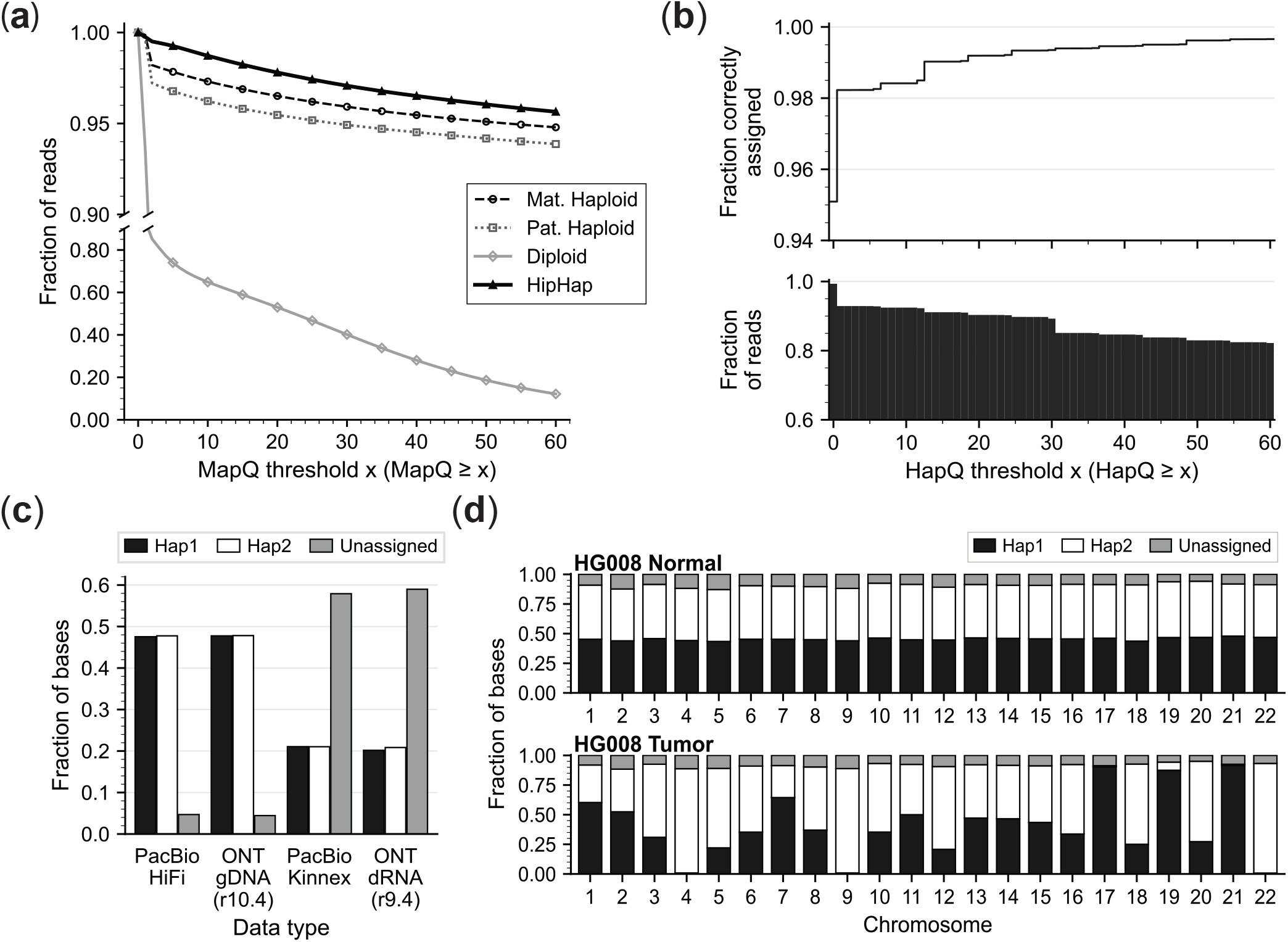
(a) Fraction of reads aligned to autosomes with MapQ at or above each threshold for four alignment strategies: maternal haploid, paternal haploid, diploid, and HipHap-processed. (b) Validation of HapQ using 200,000 simulated HG002 PacBio reads with a known haplotype of origin. (top) Fraction of all reads correctly assigned to their true haplotype, and (bottom) fraction of simulated reads at or above each HapQ score. (c) Fraction of bases in HG002 sequencing datasets assigned to each haplotype of the HG002 diploid reference assembly for diverse sequencing technologies and libraries. (d) Haplotype assignments at the chromosome level for paired HG008 normal (top) and tumor (bottom) sequencing runs aligned to the HG008N reference assembly.

To evaluate the accuracy of the calculated HapQ score, 100,000 reads were simulated from the maternal and paternal autosomes of HG002, respectively, using Badread (Wick, 2019) with the *pacbio2021* error model and a mean read length of 18*±* 5 kb. Neither junk nor random reads were included in this set. 12,821 of 198622 mapped reads had HapQ *≤* 1 and MapQ = 60, likely originating from homologous regions of the chromosomes. Conversely, 1,843 mapped reads had HapQ = 60 and MapQ *≤* 1, which were predominantly from highly repetitive centromeres and telomeres. Overall, 95.1% of reads were correctly assigned to their haplotype of origin, as shown in Figure 1b. For reads with HapQ *≥* 20 (90.3% of reads), accuracy was *∼*99.2%, increasing to *∼*99.7% for HapQ = 60 (82.1% of reads). The 0.3% mis-assignment rate at HapQ=60 (560 reads) is explained by low MapQ reads in repetitive regions of the genome where Minimap2 may miss the true assignment (253 reads) and reads that map to nearly homologous stretches of the genome with few haplotype-informative variants, resulting in HipHap marginally choosing the incorrect assignment (307 reads). The drop in reads at HapQ *>* 30, observed in Figure 1b, corresponds to a one-basepair match difference between haplotype alignments of the read under Minimap2’s `map-hifi` scoring parameters (*-*A1, -B4), leading to an AS difference of 5.

We investigated HipHap’s haplotype assignment rate across sequencing technologies, PacBio and Oxford Nanopore Technologies (ONT), and data types, genomic and transcriptomic. As shown in Figure 1c, we observed similar rates of haplotype-assigned bases in the genomic DNA reads for both the PacBio (Mat: 47.5 / Pat: 47.8 / Un: 4.7%) and ONT (Mat: 47.7 / Pat: 47.8 / Un: 4.4%) datasets. Unassigned reads were shorter than assigned reads for all datasets (Supplementary Table S1). In the transcriptomic datasets, the number of unassigned bases is greater than in genomic data for both ONT dRNA and PacBio Kinnex, because read lengths are shorter and haplotype-informative variants are less likely to occur in transcribed regions of the genome. While the rate of unassigned bases varied across all samples, the haplotype assignment rates for maternal and paternal haplotypes were nearly identical.

To validate HipHap’s ability to accurately assign haplotypes in the context of cancer-specific deletions and loss of heterozygosity, we evaluated PacBio HiFi reads of the pancreatic ductal adenocarcinoma paired tumor/normal cancer cell line HG008. HG008 provided an interesting somatic ground-truth test case, as karyotyping and careful curation by the Genome-in-a-Bottle consortium validated that numerous chromosomes exhibited complete or partial loss of heterozygosity (Wagner et al., 2026). In Figure 1d, we observed that HipHap recapitulates these events, with at most 1% of read bases mapping to each chromosome being assigned to the completely lost haplotypes on chromosomes 4, 9, 17, 21, and 22. Similarly, on chr5, where the q-arm of haplotype 1 was lost, we observe that 22% of reads were assigned to haplotype 1, 67% to haplotype 2, and 11% were unassigned (HapQ = 0). These proportions closely reflect the resolved sizes of the chr5 haplotypes in the HG008 tumor assembly: 54.2 Mbps (22.7%) of haplotype 1 and 183.7 Mbps (77.3%) of haplotype 2.

### 3.1. Runtime

To assess runtime, we evaluated both genomic and transcriptomic datasets of the HG002 cell line, aligned to the separate haplotypes of the HG002 diploid reference assembly (Hansen et al., 2025), using Minimap2 v2.30 with the parameters listed in Table 1. The mitochondrial genome assembly was included in the maternal reference.

The m21009 241011 231051 (m21009) sample was downloaded from PacBio’s public datasets (Pacific Biosciences, 2026). The PAW70337 super accuracy (sup) sample was downloaded from ONT’s 2025 Genome in a Bottle Data (GIAB) Release (Oxford Nanopore Technologies, 2025). The HG002 dRNA sample SRR30901279 was generated by Zheng *et al*. (Zheng et al., 2025). The PacBio Kinnex sample (na24385) was made publicly available by GIAB (Genome in a Bottle Consortium, 2026). The HG008 reads were generated by Wagner *et al*. (Wagner et al., 2026).

**Table 1.** Runtime benchmark datasets and parameters. Runtime was measured using 8 on an Intel Xeon Gold 6130 CPU @ 2.10 GHz.

| Table 1. Runtime benchmark datasets and parameters. Runtime was measured using 8 threads on an Intel Xeon Gold 6130 CPU @ 2.10 GHz. |  |  |  |  |  |
| --- | --- | --- | --- | --- | --- |
| Dataset | Sample ID | Number of bases (Gbps) | Minimap2 parameters | File Format | Runtime (s) |
| PacBio HiFi DNA | m21009 | 65.00 | -ax map-hifi | SAM | 1097.1 |
|  |  |  | -ax map-hifi | BAM | 1378.6 |
|  |  |  | -ax map-hifi | CRAM | 525.0 |
|  |  |  | -cx map-hifi<br>--paf-no-hit | PAF | 13.8 |
| ONT R10.4 (sup) DNA | PAW70337 | 153.68 | -ax map-ont | SAM | 1405.0 |
| ONT R9.4 direct-RNA | SRR30901279 | 75.25 | -ax splice -uf<br>-k14 | SAM | 719.1 |
| PacBio Kinnex (MasSeq) RNA | na24385 | 61.97 | -ax splice:hq<br>-uf | SAM | 648.8 |

## 4. Discussion

HipHap is an open-source tool that partitions reads between haplotypes of a diploid assembly based on independent haplotype alignments. It can be easily integrated into existing workflows between alignment and downstream analyses and works with any aligner that produces SAM, BAM, CRAM, or PAF outputs.

One limitation of the per-haplotype strategy used by HipHap is that it cannot detect structural variants that span haplotypes. For example, a read spanning a somatic fusion joining chr7 on haplotype 1 to chr11 on haplotype 2 would appear as a within-haplotype event (chr7–chr11 on haplotype 1). Although rearrangements across haplotypes are expected to be rare in germline genomes (Collins et al., 2020), they may occur in cancers, where chromosomal instability can generate complex structural variation (Li et al., 2020). To support such follow-up analyses, HipHap writes any read whose winning alignment to a haplotype spans different chromosomes to a separate fastq file. For somatic studies, we recommend re-aligning this subset of reads against the full diploid assembly to resolve putative across-haplotype events.

Several tools, such as WhatsHap, Longphase, and HiPhase (Patterson et al., 2015; Lin et al., 2022; Holt et al., 2024), can assign long reads to haplotypes using a phased variant call file (VCF). These tools parse an alignment file of reads aligned to a single haploid reference and, similarly to our HapQ proposal, tag reads with phase confidence scores (e.g., WhatsHap’s `PC:i` tag) based on the number and consistency of supporting variants.

Accurate haplotype assignment depends on both the density of heterozygous variants and the quality of the VCF input. HipHap takes a complementary approach by independently aligning reads to each haplotype assembly and comparing alignment quality directly, thereby capturing large structural differences between haplotypes that may affect alignment quality but may be absent or incompletely annotated in a VCF. Recently, a related VCF free approach was described in the Personalized Reference genome-based Cancer Genome Analysis Pipeline (PRCGAP) (Sakamoto et al., 2026), which assigns reads to a haplotype of origin using a database of haplotype-specific 21-mers built from the reference assembly. As this step is internal to the PRCGAP pipeline rather than an independent tool, we do not benchmark directly against it in this study.

Additionally, we propose HapQ as a complement to MapQ for use with diploid reference assemblies. These two scores allow users to evaluate positional (MapQ) and haplotype (HapQ) uncertainty independently, and to filter on either or both. Additional work is needed to extend this framework to polyploid genome assemblies. The HapQ concept is primarily intended for long-read alignments, as short reads typically align equally well to both haplotypes, resulting in a HapQ of 0. It may also be possible to combine the two scores into a composite score. As diploid assemblies become more widely available, we propose computing HapQ during alignment rather than as a post-processing step. This would eliminate the need to align each read set twice and address the artificial reduction in MapQ scores.

## 7. Data availability

HipHap is fully open-source and available at https://github.com/jheinz27/hiphap.

## 8. Author contributions statement

J.M.H. and H.L. conceived the project. J.M.H. implemented the method and drafted the manuscript. J.M.H. and M.G.M. analyzed the data. M.M. and H.L. provided intellectual guidance and manuscript edits. All authors read and approved the final manuscript.

## 9. Acknowledgments

We would like to thank Dr. Alaina Shumate for her valuable insights, Drs. Owen Hirschi and David Walter for their help in designing the HipHap logo, and all members of the Li and Meyerson labs for their helpful discussions.

## Appendix A: HapQ derivation

### Per-position model

Let *P*(*a, b*) be the probability of seeing a base *a* on the read and a base *b* on the haplotype at an aligned position. We always assume the joint distribution is symmetric, *P*(*a, b*) = *P*(*b, a*), and that the background base composition is uniform, *P*(*a*) = 1*/*4 for every nucleotide *a*.

### Probability of a read given a haplotype

We assume a diploid genome with two haplotypes: *h*_1_ and *h*_2_. The probability of a read sequence *r* of length *ℓ* being generated from a haplotype *h* ∈ {*h*_1_, *h*_2_} without gaps is

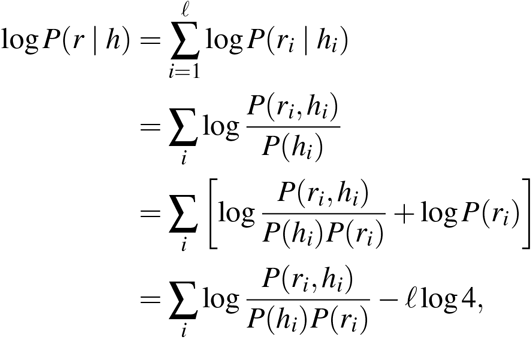

where the final line uses 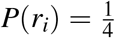, so that log *P*(*r*_*i*_) = − log 4 summed over the *l* positions. The scoring matrix *s*(·, ·) of an aligner is proportional to the per-position log-odds ratio (Durbin *et al*., 1998), so we expect

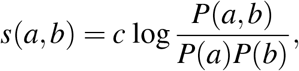

for some scaling factor *c >* 0, where *c* is defined below. The per-position sum then becomes

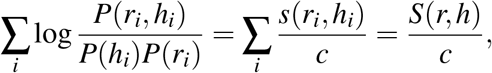

where *S*(*r, h*) = ∑_*i*_ *s*(*r*_*i*_, *h*_*i*_) is the gapless alignment score between read *r* and haplotype *h*. Hence

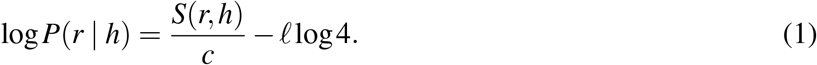

### Posterior over haplotypes

Let *h*^(*w*)^ be the sequence in the alignment of *r* against the haplotype with the better alignment score (winner) and *h*^(*l*)^ be the sequence of the other haplotype (loser). When the alignment scores to the two haplotypes are equal, *h*^(*w*)^ and *h*^(*l*)^ are randomly assigned. Without loss of generality, suppose *P*(*h*^(*w*)^ | *r*) ≥ *P*(*h*^(*l*)^ | *r*). Under a flat prior *P*(*h*^(*w*)^) = *P*(*h*^(*l*)^) = 0.5, we can compute the posterior:

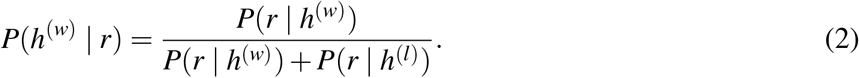

### From posterior to HapQ

The HapQ score is the Phred-scaled posterior probability that the winning assignment is incorrect,

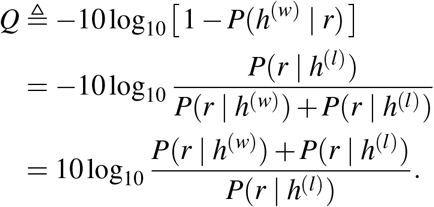

where the second equality uses 1 − *P*(*h*^(*w*)^ |*r*) = *P*(*h*^(*l*)^ |*r*). On the condition *P*(*h*^(*w*)^ |*r*) ≫*P*(*h*^(*l*)^ |*r*), the losing term is negligible in the numerator, so that *P*(*r*| *h*^(*w*)^) + *P*(*r*| *h*^(*l*)^) ≈*P*(*r* |*h*^(*w*)^); this condition is easily met in practice, since the likelihood ratio is exponential in the alignment-score gap, so even a small score difference makes *P*(*r*| *h*^(*l*)^) negligible. Substituting the log-probability relation in eq. 1 and applying this approximation,

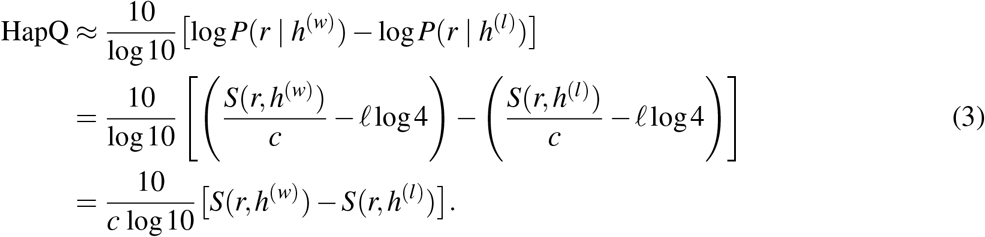

### Determining the scaling constant *c*

It remains to fix *c*. This is achieved by evaluating the score at a perfect match case (*a* = *b*), which is the per-base match score *s*(*a, a*) used by the alignment software, with per-base error rate *ε*,

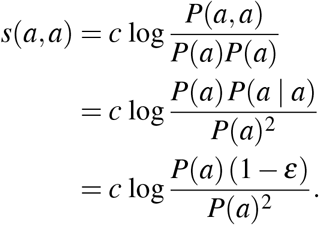

Under a low error rate, 1 − *ε* ≈ 1.

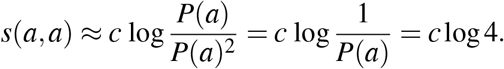

We therefore define the scaling constant so that a perfect match maps exactly to log 4,

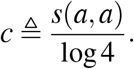

Substituting this *c* into eq. 3 gives the final HapQ,

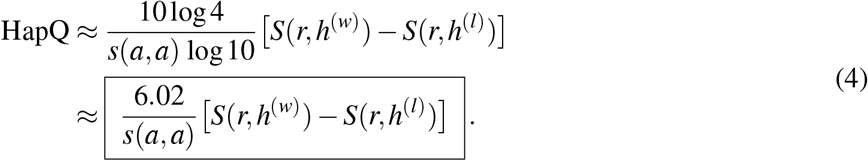

The HapQ framework can be extended to gap penalties if they are on a similar scale as mismatches.

### Supplementary Tables

**Supplementary Table S1:** Read counts and read-length summary statistics stratified by HapQ. For each sample, the number of reads and the mean and standard deviation of read length (bp) are reported separately for haplotype assigned reads (HapQ ≥ 2) and unassigned reads (HapQ ≤ 1).

| Sample | HapQ $\geq 2$ | | | HapQ $\leq 1$ | | |
| --- | --- | --- | --- | --- | --- | --- |
|  | # reads | Mean | Std | # reads | Mean | Std |
| HG002_PB_gDNA | 3,076,836 | 20,165.9 | 6,328.4 | 183,984 | 15,988.7 | 7,052.2 |
| HG002_ONT_gDNA | 6,960,273 | 21,105.6 | 15,559.1 | 1,242,149 | 5,289.5 | 8,329.0 |
| HG002_ONT_dRNA | 29,142,063 | 1,217.7 | 957.8 | 42,779,397 | 926.5 | 720.2 |
| HG002_PB_MasSeq_RNA | 20,497,757 | 1,401.9 | 619.8 | 26,654,550 | 1,243.5 | 519.6 |
| HG008N_PB_gDNA | 5,969,711 | 16,516.6 | 4,105.7 | 709,504 | 15,561.5 | 3,837.8 |
| HG008T_PB_gDNA | 12,665,361 | 17,616.1 | 4,318.8 | 1,348,766 | 16,569.2 | 4,014.4 |

## References

J. K. Bonfield, J. Marshall, P. Danecek, H. Li, and V. Ohan, et al. HTSlib: C library for reading/writing high-throughput sequencing data. GigaScience, 10(2):giab007, Feb. 2021. ISSN 2047-217X. doi: 10.1093/gigascience/giab007.

H. Cheng, G. T. Concepcion, X. Feng, H. Zhang, and H. Li. Haplotype-resolved de novo assembly using phased assembly graphs with hifiasm. Nature Methods, 18(2):170–175, Feb. 2021. ISSN 1548-7105. doi: 10.1038/s41592-020-01056-5.

H. Cheng, H. Qu, S. McKenzie, K. R. Lawrence, and R. Windsor, et al. Efficient near-telomere-to-telomere assembly of nanopore simplex reads. Nature, pages 1–8, Feb. 2026. ISSN 1476-4687. doi: 10.1038/s41586-026-10105-6.

R. L. Collins, H. Brand, K. J. Karczewski, X. Zhao, and J. Alföldi, et al. A structural variation reference for medical and population genetics. Nature, 581(7809):444–451, May 2020. ISSN 1476-4687. doi: 10.1038/s41586-020-2287-8.

R. Durbin, S. R. Eddy, A. Krogh, and G. Mitchison. Biological Sequence Analysis: Probabilistic Models of Proteins and Nucleic Acids. Cambridge University Press, Cambridge, UK, 1998. ISBN 978-0-521-54079-7.

S. Garg, M. Rautiainen, A. M. Novak, E. Garrison, and R. Durbin, et al. A graph-based approach to diploid genome assembly. Bioinformatics, 34(13):i105–i114, July 2018. ISSN 1367-4803. doi: 10.1093/bioinformatics/bty279.

Genome in a Bottle Consortium. PacBio MAS-Seq (Kinnex) FLNC alignments for HG002/NA24385. https://ftp-trace.ncbi.nlm.nih.gov/giab/ftp/data_RNAseq/AshkenazimTrio/HG002_NA24385_son/PacBio_Pacbio-MASseq/GM24385/2-FLNC/, 2026. Accessed 17 July 2026.

N. F. Hansen, N. Dwarshuis, H. J. Ji, A. Rhie, and H. Loucks, et al. A complete diploid human genome benchmark for personalized genomics. bioRxiv, page 2025.09.21.677443, Sept. 2025. ISSN 2692-8205. doi: 10.1101/2025.09.21.677443.

J. M. Heinz, M. Meyerson, and H. Li. Detecting foldback artifacts in long-reads. BMC Genomics, 27(1):144, Jan. 2026. ISSN 1471-2164. doi: 10.1186/s12864-025-12492-y.

J. M. Holt, C. T. Saunders, W. J. Rowell, Z. Kronenberg, and A. M. Wenger, et al. HiPhase: Jointly phasing small, structural, and tandem repeat variants from HiFi sequencing. Bioinformatics, 40(2):btae042, Feb. 2024. ISSN 1367-4811. doi: 10.1093/bioinformatics/btae042.

International Human Genome Sequencing Consortium. Initial sequencing and analysis of the human genome. Nature, 409 (6822):860–921, Feb. 2001. ISSN 1476-4687. doi: 10.1038/35057062.

A. G. Keskus, A. Bryant, T. Ahmad, B. Yoo, and S. Aganezov, et al. Severus detects somatic structural variation and complex rearrangements in cancer genomes using long-read sequencing. Nature Biotechnology, 44(2):247–257, Feb. 2026. ISSN 1546-1696. doi: 10.1038/s41587-025-02618-8.

J. Köster. Rust-Bio: A fast and safe bioinformatics library. Bioinformatics, 32(3):444–446, Feb. 2016. ISSN 1367-4803. doi: 10.1093/bioinformatics/btv573.

H. Li. Aligning sequence reads, clone sequences and assembly contigs with BWA-MEM, May 2013.

H. Li. Minimap2: Pairwise alignment for nucleotide sequences. Bioinformatics, 34(18):3094–3100, Sept. 2018. ISSN 1367-4803. doi: 10.1093/bioinformatics/bty191.

H. Li, B. Handsaker, A. Wysoker, T. Fennell, and J. Ruan, et al. The Sequence Alignment/Map format and SAMtools. Bioinformatics, 25(16):2078–2079, Aug. 2009. ISSN 1367-4803. doi: 10.1093/bioinformatics/btp352.

Y. Li, N. D. Roberts, J. A. Wala, O. Shapira, and S. E. Schumacher, et al. Patterns of somatic structural variation in human cancer genomes. Nature, 578(7793):112–121, Feb. 2020. ISSN 1476-4687. doi: 10.1038/s41586-019-1913-9.

W.-W. Liao, M. Asri, J. Ebler, D. Doerr, and M. Haukness, et al. A draft human pangenome reference. Nature, 617 (7960):312–324, May 2023. ISSN 1476-4687. doi: 10.1038/s41586-023-05896-x.

J.-H. Lin, L.-C. Chen, S.-C. Yu, and Y.-T. Huang. LongPhase: An ultra-fast chromosome-scale phasing algorithm for small and large variants. Bioinformatics, 38(7):1816–1822, Mar. 2022. ISSN 1367-4803. doi: 10.1093/bioinformatics/btac058.

S. Nurk, S. Koren, A. Rhie, M. Rautiainen, and A. V. Bzikadze, et al. The complete sequence of a human genome. Science, 376(6588):44–53, Apr. 2022. doi: 10.1126/science.abj6987.

Oxford Nanopore Technologies. Genome in a Bottle Data Release 2025.01. https://epi2me.nanoporetech.com/giab-2025.01/, 2025. Accessed 17 July 2026.

Pacific Biosciences. PacBio-Datasets-Highly accurate long-read sequencing. https://www.pacb.com/connect/datasets/, 2026. Accessed 17 July 2026.

M. Patterson, T. Marschall, N. Pisanti, L. van Iersel, and L. Stougie, et al. WhatsHap: Weighted Haplotype Assembly for Future-Generation Sequencing Reads. Journal of Computational Biology, 22(6):498–509, June 2015. ISSN 1066-5277. doi: 10.1089/cmb.2014.0157.

M. Rautiainen, S. Nurk, B. P. Walenz, G. A. Logsdon, and D. Porubsky, et al. Telomere-to-telomere assembly of diploid chromosomes with Verkko. Nature Biotechnology, 41(10): 1474–1482, Oct. 2023. ISSN 1546-1696. doi: 10.1038/s41587-023-01662-6.

Y. Sakamoto, Y. Ochi, Y. Kogure, S. Kato, and A. Sato-Otsubo, et al. Personalized reference genome-based pipeline reveals comprehensive haplotype-resolved views of cancer genomes, July 2026. ISSN 2692-8205.

V. A. Schneider, T. Graves-Lindsay, K. Howe, N. Bouk, and H.-C. Chen, et al. Evaluation of GRCh38 and de novo haploid genome assemblies demonstrates the enduring quality of the reference assembly. Genome Research, 27(5):849–864, May 2017. ISSN 1549-5469. doi: 10.1101/gr.213611.116.

M. Smolka, L. F. Paulin, C. M. Grochowski, D. W. Horner, and M. Mahmoud, et al. Detection of mosaic and population-level structural variants with Sniffles2.Nature Biotechnology, 42 (10):1571–1580, Oct. 2024. ISSN 1546-1696. doi: 10.1038/s41587-023-02024-y.

J. Wagner, A. G. Keskus, K. K. Oshima, T. R. Ranallo-Benavidez, and J. McDaniel, et al. A complete human pancreatic cancer genome. bioRxiv, page 2026.05.01.722316, Jan. 2026. doi: 10.64898/2026.05.01.722316.

R. R. Wick. Badread: Simulation of error-prone long reads. Journal of Open Source Software, 4(36):1316, Apr. 2019. ISSN 2475-9066. doi: 10.21105/joss.01316.

Z. Zheng, X. Yu, L. Chen, Y.-L. Lee, and C. Xin, et al. Clair3-RNA: A deep learning-based small variant caller for long-read RNA sequencing data. Nature Communications, 16(1):11553, Dec. 2025. ISSN 2041-1723. doi: 10.1038/s41467-025-67237-y.

